# AI-enhanced cardiac digital twins extend drug proarrhythmic risk assessment through experimental data uncertainty propagation and overdose exploration: a loperamide case study

**DOI:** 10.64898/2026.01.28.702200

**Authors:** Paula Dominguez-Gomez, Alberto Zingaro, Caterina Balzotti, Michael Leitner, Noelia Jacobo-Piqueras, Mariano Vazquez, Georg Rast, Jazmin Aguado-Sierra

## Abstract

Drug-induced QT interval prolongation is a key biomarker of proarrhythmic risk and central to drug cardiac safety evaluation alongside in vitro assays and animal studies. Current preclinical frameworks, however, provide limited insight into how experimental uncertainty and extreme exposures translate into real-world arrhythmic risk, despite both factors critically modulating outcomes.

To address this, we used sex-specific machine learning surrogate models trained on 3D cardiac digital twins—mechanistic electrophysiology models of anatomically detailed ventricles that integrate multichannel ion-channel block data. These emulators combine the realism of 3D simulations with high-throughput capability, enabling rapid, ethically unconstrained assessment of proarrhythmic risk.

We illustrate the approach using loperamide, safe at therapeutic doses but linked to fatal arrhythmias at extreme exposures. Two analyses were performed: propagating experimental IC_50_ and Hill coefficient variability to quantify its effect on predicted QT prolongation and arrhythmic probability, and simulating extreme exposures to identify sex-specific arrhythmogenic thresholds.

Experimental variability substantially broadened predicted QT prolongation and arrhythmic risk near decision thresholds. Extreme exposure simulations identified arrhythmogenic thresholds of approximately 107–109 × *C*_max_ in female models and 213–286 × *C*_max_ in male models. This framework offers a scalable, physics-based tool for early-stage drug cardiac safety evaluation.

## 1 Introduction

Drug-induced QT interval prolongation (ΔQT) is a well-established surrogate marker of proarrhythmic risk, particularly torsades de pointes, and remains a leading cause of post-approval restrictions or withdrawals (Shah, 2002). Existing safety assessment frameworks, comprising in vitro ion-channel assays, in vivo animal studies, and clinical thorough-QT (TQT) trials, are effective at identifying hazards within expected therapeutic exposure ranges. However, they are not designed to systematically address conditions such as deliberate overdose, drug–drug interactions, or metabolic impairment, where plasma concentrations can greatly exceed therapeutic levels. These extreme exposure scenarios are rarely evaluated before marketing, despite their potential to reveal serious arrhythmic liabilities (Vargas et al., 2021).

Computational modeling has emerged as a powerful complement to experimental testing in this context. Mechanistic cardiac electrophysiology models can integrate multichannel ion-block data with physiological detail to predict drug effects on repolarization and arrhythmic risk (Trayanova et al., 2024; Goldring et al., 2025). These models enable in silico exploration of scenarios that are impractical or unethical to test in humans, including very high drug concentrations and complex ion-channel interaction patterns (Mirams et al., 2011; Wilhelms et al., 2012; Zemzemi et al., 2013; Passini et al., 2017; Abbasi et al., 2017; Sahli Costabal et al., 2018; Okada et al., 2018; Romero et al., 2018; Passini et al., 2019; Li et al., 2019; Hwang et al., 2019; Llopis-Lorente et al., 2023; Aguado-Sierra et al., 2024). Importantly, three-dimensional (3D) cardiac models provide anatomically and electrophysiologically realistic representations that capture spatial heterogeneity of activation and repolarization, enabling direct computation of pseudo-ECGs and ΔQT. Previous work has shown that sex-specific 3D models capture complex drug-induced proarrhythmic mechanisms that simpler models can not (Dominguez-Gomez et al., 2024), highlighting their importance for translational safety studies.

Yet, the predictive value of such simulations depends critically on the quality and variability of their inputs. Experimental measurements of ion-channel blockade, such as IC_50_ values, Hill coefficients, and binding kinetics, can vary across laboratories and assays (Elkins et al., 2013). This variability propagates through the modeling chain, influencing predicted ΔQT and arrhythmia likelihood. However, the computational cost of high-fidelity 3D simulations has historically prevented their use for exhaustive uncertainty quantification or large-scale dose–response exploration (Bridio et al., 2025).

In our previous work (Dominguez-Gomez et al., 2024), we addressed this computational bottleneck by introducing machine learning–based surrogate models (emulators) trained on sex-specific 3D cardiac electrophysiology digital twins. These are surrogate models that learn the input–output mapping between ion-channel block profiles, drug concentration, and relevant biomarkers such as ΔQT and arrhythmia incidence (Sahli Costabal et al., 2019; Grandits et al., 2024; Dominguez-Gomez et al., 2024), while reproducing the predictive accuracy of high-fidelity simulations at a fraction of the computational cost. This advance enables rapid and reproducible exploration of large parameter spaces that would otherwise be computationally prohibitive.

Crucially, these emulators provide a general framework to assess any drug for which ion-channel data are available. They are particularly well suited to address two major challenges in modern safety pharmacology: quantifying the impact of experimental data variability on predicted outcomes, and systematically exploring supratherapeutic and extreme exposure scenarios that lie outside standard regulatory testing paradigms. The sex-specific nature of these models is particularly relevant given growing evidence that drug-induced proarrhythmic risk can differ substantially between males and females due to electrophysiological, hormonal, and structural factors (Darpo et al., 2014; Peirlinck et al., 2021, Peirlinck et al., 2022; Dominguez-Gomez et al., 2025). Loperamide, an over-the-counter antidiarrheal that acts as a peripheral *µ*-opioid receptor agonist, provides an illustrative test case for such a framework (DrugBank, 2025). At therapeutic doses, it is generally considered safe and is excluded from central nervous system effects due to poor blood–brain barrier penetration.

However, in recent years, loperamide has been associated with intentional misuse and abuse, where individuals ingest massive doses in attempts to self-treat opioid withdrawal or achieve euphoric effects (Vakkalanka et al., 2017). At these supratherapeutic concentrations, loperamide can cross the blood–brain barrier and also inhibit multiple cardiac ion channels. Such multichannel block at extreme exposures has been linked to serious ventricular arrhythmias and sudden cardiac death in case reports and pharmacovigilance data (Lee et al., 2019).

In this study, building directly on our previously developed emulators, we investigate how experimental ion-channel variability and extreme drug exposure jointly shape the concentration–ΔQT relationship and proarrhythmic risk profile of loperamide. By propagating experimentally observed variability in ion-channel block parameters through sex-specific AI-enhanced cardiac emulators, we perform large-scale uncertainty quantification across both therapeutic and overdose concentration ranges. This approach enables rapid identification of sex-dependent thresholds for QT prolongation and arrhythmic onset. To our knowledge, this is the first study to combine sex-specific 3D cardiac digital twins, machine-learning emulation, and systematic quantification of experimental ion-channel uncertainty to assess proarrhythmic risk under extreme drug exposure.

## 2 Methodology

The complete methodology implemented in this study is summarized in Figure 1. Starting from experimental ion-channel block data measured under different conditions (saline buffer and human plasma) (Section 2.1), variability in IC_50_ values and Hill coefficients is explicitly sampled to generate probabilistic input distributions (Section 2.2). These sampled parameters are combined with drug concentration using a Hill-type model to compute concentration-dependent block of seven major cardiac ionic currents. The resulting multichannel block profiles are provided as inputs to sex-specific AI-enhanced emulators trained on a high-fidelity three-dimensional cardiac digital twin or simulator (Section 2.3). These emulators rapidly predict ΔQT and arrhythmic outcomes while preserving the mechanistic fidelity of the underlying 3D simulator across both therapeutic and extreme exposure scenarios (Section 2.4). By propagating input uncertainty through the emulators, the framework enables fast uncertainty quantification of ΔQT and proarrhythmic risk.

**Figure 1:**
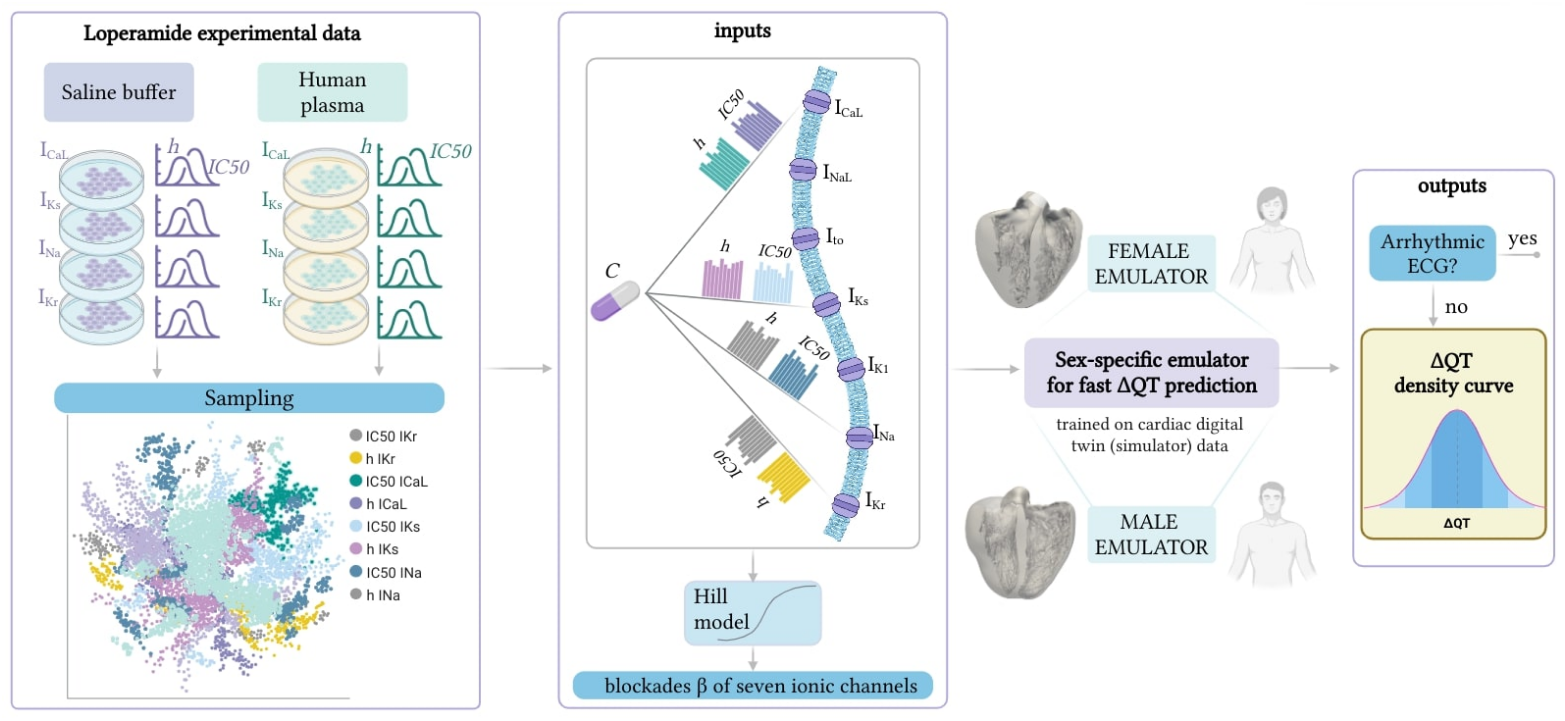
Overview of the proposed methodology. The workflow links experimental ion-channel data to predictions of ΔQT and arrhythmic risk. Experimental measurements are sampled within their reported variability to capture the uncertainty space. This sampled data is translated into ion-channel blocks that serve as inputs for the sex-specific emulators.

### 2.1 Loperamide data

Loperamide is an over-the-counter antidiarrheal agent that acts as a peripheral *µ*-opioid receptor agonist (Akel and Bekheit, 2018). At therapeutic doses, it is generally considered safe; however, in cases of misuse or overdose, it has been associated with inhibition of multiple cardiac ion channels and subsequent arrhythmic events (Marraffa et al., 2014).

Experimental data characterizing loperamide’s ion channel profile was generated by in vitro patch-clamp experiments (Table 1). Blockade was quantified for the most affected ion channels: *I*_Kr_, *I*_CaL_, *I*_Na_, and *I*_Ks_. For each channel, concentration–response curves were obtained from replicate experiments using cells expressing the corresponding ion channel, from which IC_50_ values and Hill coefficients were derived. In the case of hERG, experiments were repeated in two different media—saline buffer and 100% human plasma—to capture protein-binding effects, whereas for the other channels only saline buffer conditions were tested. The intrinsic variability in the input data was characterized from the replicate measurements.

**Table 1:**
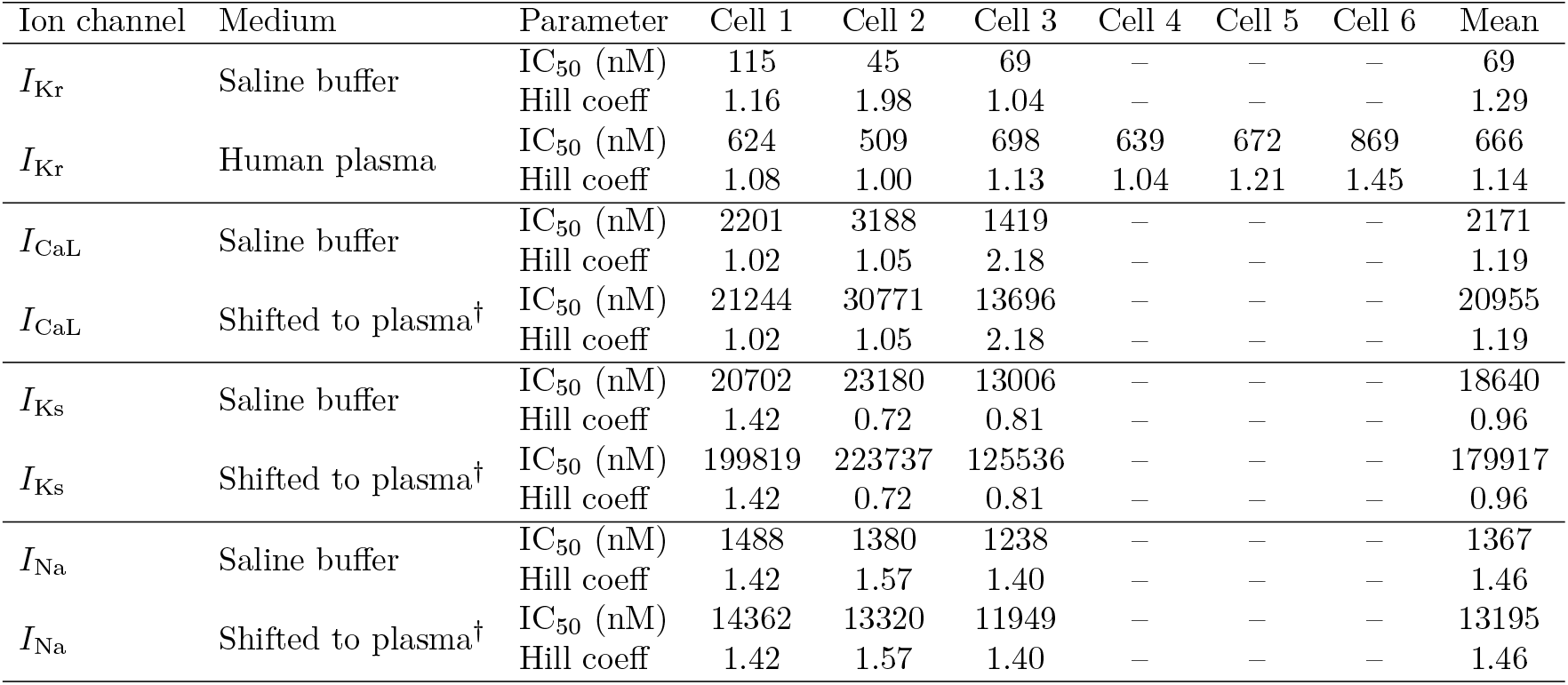
Loperamide ion-channel inhibition parameters obtained from in vitro assays. Each column labelled Cell 1–6 corresponds to an independent replicate experiment performed using cells expressing the respective ion channel. Mean values across replicates are reported in the rightmost column. For hERG (*I*_Kr_), experiments were performed in both saline buffer and 100% human plasma to evaluate protein-binding effects. ^†^ Non-hERG channels were only measured in saline; IC_50_ values were shifted to plasma using the hERG-derived saline-to-plasma factor of 9.652 and Hill coefficients were kept unchanged.

Because only a fraction of the drug circulating is pharmacologically active, translating saline measurements to in vivo–relevant conditions requires accounting for functional plasma protein binding (functional PPB).

Functional PPB reflects the effective fraction of free drug available to block ion channels under physiological conditions (Smith et al., 2010). In the hERG assays, the comparison between saline and plasma revealed a 9.65-fold rightward shift in IC_50_, corresponding to a functional PPB of 89.64%. Published PPB values for loperamide (95%, reported in DrugBank (2025)), however, differ from this estimate, underscoring that functional shifts determined experimentally may not align with calculated PPB values. Since *I*_CaL_, *I*_Ks_, and *I*_Na_ were measured only in saline, these channels were adjusted by applying the same experimentally determined shift for *I*_Kr_. This correction ensures consistency across channels and avoids underestimating their contribution to loperamide’s multichannel block. Importantly, functional plasma protein binding is not routinely measured in safety pharmacology, yet our results highlight its critical role in producing physiologically meaningful input data for in silico modeling.

### 2.2 Uncertainty propagation analysis

#### 2.2.1 Combinatorial sampling

Ion-channel blockade data are inherently variable due to interexperimental differences. To explicitly account for this variability, we propagated uncertainty by considering all available experimental replicates of IC_50_ and Hill coefficients. For each medium, we systematically combined values across channels to generate comprehensive input sets. In saline buffer, three replicates were available per channel (Table 1), which resulted in 3^4^ = 81 unique combinations of IC_50_ and Hill parameter sets. In human plasma, six replicates were available for hERG and three for the remaining channels, yielding a total of 6 × 3^3^ = 162 combinations. Each of these parameter sets was used as input to the emulators to predict concentration-ΔQT and arrhythmic outcome across the exposure range. This procedure corresponds to an exhaustive combinatorial (full factorial) sampling of replicates without replacement (Saltelli et al., 2008), ensuring that all possible combinations of experimental inputs are explored. Such an approach captures the full variability envelope of experimental data; however, it does so in a discrete and non-homogeneous manner, potentially overrepresenting specific values.

#### 2.2.2 Sobol sequences

To obtain a more uniform and continuous coverage of the uncertainty space, the analysis was complemented with a quasi-random sampling approach based on Sobol sequences. Sobol sampling provides low-discrepancy, quasi–Monte Carlo points that fill multidimensional spaces more evenly than purely random draws, improving convergence and representativeness in nonlinear systems such as cardiac electrophysiology models (Saltelli et al., 2008; Renardy et al., 2021). To apply this method, the replicate IC_50_ and Hill coefficient values for each ion channel were used to compute their mean (*µ*) and standard deviation (*σ*), from which sampling ranges are defined as [*µ −* 1.1*σ, µ* + 1.1*σ*]. These bounds correspond approximately to the central 75% of a Gaussian distribution, which was assumed to describe the data if a larger number of measurements were available. The slight expansion to *±*1.1*σ* ensured that marginal experimental values remain represented while excluding unrealistic extremes. This design balances realism and variability, providing a statistically homogeneous sampling of the uncertainty space and enabling more efficient exploration of input–output relationships.

### 2.3 Computational framework

Previously developed and validated sex-specific cardiac emulators (Dominguez-Gomez et al., 2024) were employed. These emulators act as fast approximations of a high-fidelity simulator (Aguado-Sierra et al., 2024), enabling rapid prediction of drug-induced QT prolongation. Briefly, the underlying simulator relies on Alya, a high-performance finite-element framework for cardiac electrophysiology (Vázquez et al., 2016; Santiago et al., 2018) that solves the monodomain equations of cardiac electrophysiology coupled with the O’Hara–Rudy cellular model, extended with sex-specific ionic conductances (O’Hara et al., 2011; Passini et al., 2016; Iseppe et al., 2021).

Drug effects are modeled via a multichannel conductance block formulation (Mirams et al., 2011). Given the conductance *g*_k_ of one of the seven most influent ionic channels *I*_CaL_, *I*_NaL_, *I*_to_, *I*_Ks_, *I*_K1_, *I*_Na_, and *I*_Kr_ Crumb et al. (2016), we define the ion channel conductance after the drug administration with the following Hill model:

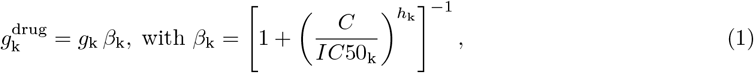

with k = CaL, NaL, to, Ks, K1, Na, Kr. In the equation above, 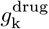 is the conductance of the k–th channel after drug administration. Thus, the blockade *β*_k_ of channel k is identified by the Hill coefficient *h*_k_, the *IC*50_k_ and the drug concentration *C*. Notice that *C* is defined as the concentration of the drug in the plasma that is not bound to plasma proteins and is therefore available to exert a pharmacological effect.

These are the components that physiologically reproduce the activation and repolarization across anatomically detailed biventricular meshes, from which pseudo-ECGs and ΔQT can be directly computed (Gonzalez-Martin et al., 2023). While this simulator provides physiologically realistic predictions of drug effects on the QT interval, its high computational cost prevents high-throughput exploration.

The emulators used in this work were trained on a large dataset of 900 high-fidelity 3D simulations (450 per sex), spanning a wide range of multichannel block profiles (Dominguez-Gomez et al., 2024). The architecture of the emulators integrates a classifier followed by a regressor. The classifier enhances the robustness and accuracy of our emulators by filtering out non-arrhythmic and arrhythmic cases, reducing dataset complexity and discontinuity, while the regressor predicts ΔQT values. Therefore, the inputs to this modeling pipeline are the fractional blocks of the ionic currents affected by the drug of study, and their outputs are ΔQT values together with the occurrence or absence of arrhythmic events.

These emulators reproduce simulator’s outputs with *<*4% error while reducing computational cost by five orders of magnitude, making them essential for uncertainty quantification in this study. Importantly, this methodology has been validated against clinical data, where emulator-based predictions were shown to agree with observed concentration–ΔQT relationships across multiple compounds. A more detailed description of this methodology can be found in (Dominguez-Gomez et al., 2024).

### 2.4 Extreme exposure analysis

To evaluate the potential for arrhythmia under conditions of deliberate overdose or misuse, the simulations were extended far beyond therapeutic exposure. The mean total therapeutic plasma concentration (C_max_) of loperamide, reported as 3.98 ng/mL (Lu et al., 2023), was used as the reference concentration. Following a strategy similar to that described by Lu et al. (2023), a concentration–response protocol spanning from 1×C_max_ to 2000×C_max_ was constructed. A total of 208 concentrations were sampled across this range using a uniform distribution. The intention of this extreme exposure scenario was to define safety margins by identifying the highest exposure associated with stable repolarization and to locate arrhythmic thresholds that could then be compared against concentrations reported in clinical case studies of loperamide overdose. Moreover, for every given concentration, the probability of arrhythmia (*P* (Arrhythmia)) was computed as the fraction of sampled simulations classified as arrhythmic separately for each sex and sampling strategy. This approach enables a systematic assessment of whether the predicted onset of arrhythmia coincides with observed adverse outcomes, thereby strengthening the translational value of the computational framework.

### 2.5 Targeted emulators’ validation under extreme conditions

Although the emulators were previously validated against a set of benchmark drugs (Dominguez-Gomez et al., 2024), their application here extends beyond the conditions explicitly covered in that prior work. In particular, the present analysis explores (i) compound-specific ion-channel uncertainty derived from replicate loperamide experiments and (ii) extreme exposure regimes approaching the onset of arrhythmic instability. Both aspects probe regions of the input space where nonlinear dynamics and classification boundaries may challenge surrogate-model accuracy. While the emulators were trained to reproduce simulator outputs across a broad range of conditions, their predictive accuracy is expected to decrease near arrhythmic transitions, particularly under extreme ion channel blockades that are sparsely represented in the training data.

This limitation is intrinsic to surrogate modeling and for this reason a targeted validation was performed to ensure that the emulators faithfully reproduce the simulator’s behavior across both safe and arrhythmogenic regimes, confirming that their predictive accuracy extends to the compound- and exposure-dependent scenarios analyzed here. Saline buffer data was used for this purpose.

Validation scenarios were identified based on the methods described in Sections 2.2 and 2.4. From the ensemble of sampled ion-channel parameter combinations, we selected two configurations that maximized divergence in predicted outcomes. Divergence was defined as the largest separation in predicted ΔQT at identical concentrations. These two configurations were then simulated in both male and female 3D cardiac digital twins at three representative concentrations of loperamide: 82 ng/mL, 1999 ng/mL, and 4033 ng/mL. The first scenario, at 82 ng/mL, corresponds to a clearly non-arrhythmic supratherapeutic level roughly twenty-fold higher than the reported therapeutic C_max_ of 3.98 ng/mL. The second scenario, at 1999 ng/mL, represents an intermediate regime near the threshold of arrhythmic onset identified in the emulator-based analysis, while the third scenario, at 4033 ng/mL, reflects extreme exposures that would be expected to produce severe repolarization disturbances and torsades de pointes.

The purpose of this selection was to challenge the emulators across the full spectrum of behaviors relevant to proarrhythmic risk assessment, from stable QT prolongation to overt arrhythmia. For each scenario, emulators’ predictions of ΔQT and arrhythmic outcome were directly compared against results obtained from the simulator.

## 3 Results

This section presents the results of the emulator-based analyses, structured to elucidate how experimental variability and extreme exposure shape proarrhythmic risk and build confidence in the predictions. We first characterize the concentration–ΔQT relationship under propagated ion-channel uncertainty using the two sampling strategies (Section 3.1). We then quantify the incidence of arrhythmic events and identify sex-specific exposure thresholds at which arrhythmias emerge (Section 3.2). Finally, to ensure the reliability of the emulator-based findings in the most critical regions of the parameter space, we directly compare emulator predictions against high-fidelity simulations (Section 3.3).

### 3.1 Concentration–response patterns

To evaluate how experimental uncertainty influences ΔQT predictions of the emulators, the results obtained from both sampling strategies were analyzed: the discrete combinatorial sampling of experimental replicates and the quasi-random Sobol sequence sampling based on statistically defined parameter ranges. In both analyses, predictions classified as arrhythmic were excluded from the ΔQT analysis, as the QT interval can not be well defined once stable repolarization is lost.

#### 3.1.1 Combinatorial sampling

Figure 2 summarizes the relationship between loperamide concentration and ΔQT across both media conditions. At the broadest exposure scale (Figures 2A–2B), both saline buffer and human plasma data show a steep nonlinear increase in ΔQT with concentration. Zoomed-in views of the therapeutic range reveal negligible changes in ΔQT, consistent with the lack of cardiac risk under therapeutic dosing.

**Figure 2:**
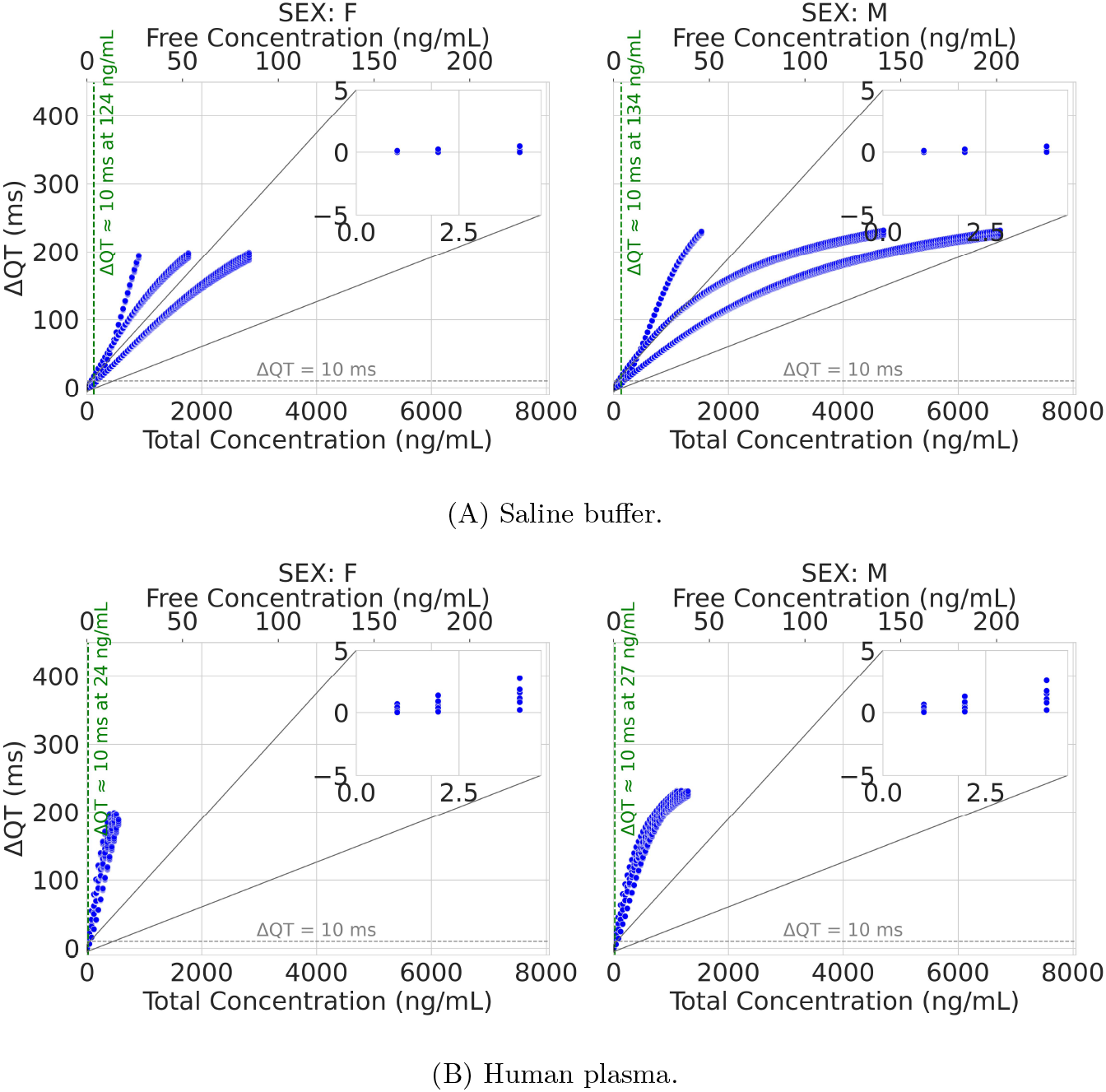
Emulators’ predictions of ΔQT as a function of loperamide concentration for female (left) and male (right) using combinatorial sampling. Main panels show the full concentration range, with insets highlighting the therapeutic exposure range. The bottom x-axis reports free (unbound fraction responsible for pharmacological activity) plasma concentration and the top x-axis the corresponding total concentration (bound + unbound); both are linked via plasma protein binding and represent the same simulations. Arrhythmic predictions were excluded, as ΔQT is not computable once arrhythmic dynamics occur.

Variability in ΔQT predictions is already visible at low supratherapeutic concentrations particularly for the saline buffer data, with predictions diverging by tens of milliseconds at identical nominal doses; this dispersion increased with concentration. Concentrations at which the cohort mean ΔQT exceeded 10 ms occur well above therapeutic levels. Importantly, arrhythmic outcomes in these simulations are associated with substantially larger ΔQT values (typically on the order of hundreds of milliseconds). Comparisons between saline and plasma-adjusted datasets reveal that the plasma condition consistently narrowed the spread of ΔQT predictions, reflecting the dampening effect of protein binding on effective drug potency. This reduction in variability suggests that saline-based measurements may overestimate the uncertainty of in vivo responses, while plasma-adjusted data provide a more constrained and physiologically relevant distribution.

The combinatorial approach captures all extreme combinations of input parameters, thereby delineating the full experimental uncertainty envelope; however, the resulting distribution is discrete and sometimes uneven, with dense clustering of points corresponding to specific replicate combinations.

#### 3.1.2 Sobol sequences

To complement the discrete replicate-based analysis, a quasi-random sampling based on Sobol sequences was performed to obtain a smoother and more homogeneous exploration of the uncertainty space. Figure 3 shows emulators-predicted ΔQT as a function of concentration. In both media, the Sobol sampling reproduces the nonlinear concentration-response relationship observed with the combinatorial analysis but distributes outcomes more uniformly across the parameter space.

**Figure 3:**
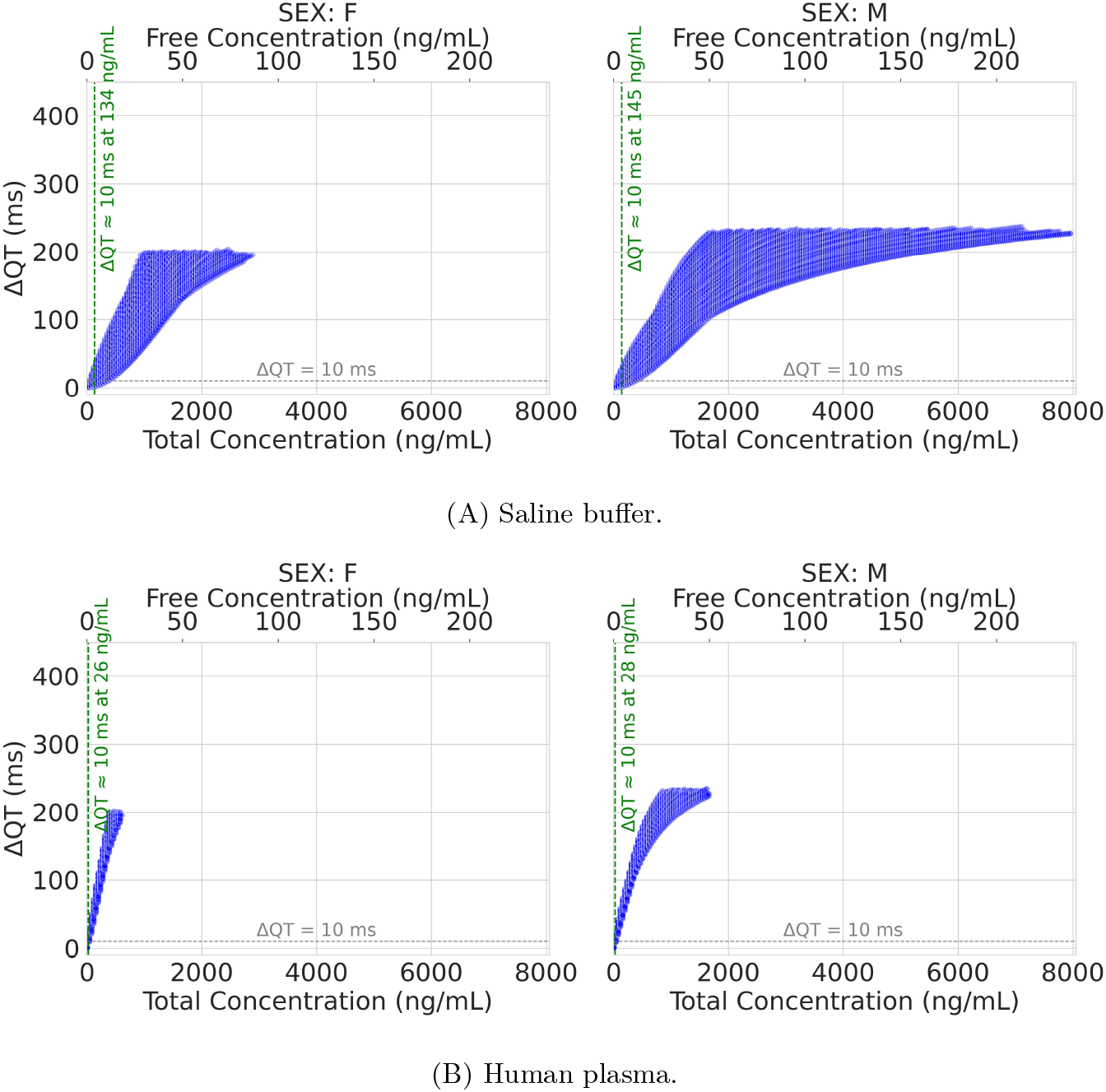
Emulators’ predictions of ΔQT as a function of loperamide concentration for female (left) and male (right) using sobol sampling. The bottom x-axis reports free (unbound fraction responsible for pharmacological activity) plasma concentration and the top x-axis the corresponding total concentration (bound + unbound); both are linked via plasma protein binding and represent the same simulations. Arrhythmic predictions were excluded, as ΔQT is not computable once arrhythmic dynamics occur.

In saline buffer (Figure 3A), variability among virtual individuals remains high near the arrhythmic threshold (the concentration at which the first arrhythmia appears), and becomes progressively constrained beyond this point, as most simulations are classified as arrhythmic and excluded from the analysis. The plasma-adjusted condition (Figure 3B) produces higher overall ΔQT values and advances arrhythmic onset. The concentration corresponding to a mean ΔQT of 10 ms occurrs at 134 ng/mL in females and 145 ng/mL in males for plasma data, compared with 26 ng/mL for females and 28 ng/mL for males for saline data, in line with combinatorial sampling predictions.

Because Sobol sequences ensure uniform coverage of the multidimensional uncertainty space, the resulting distributions avoid the artificial clustering seen in the combinatorial design. Variability remains the largest near the transition zone between stable repolarization and arrhythmic onset, reinforcing that small parameter perturbations in this region can toggle outcomes between safe and arrhythmogenic states. These results confirm that Sobol sampling provides a statistically homogeneous and computationally efficient means of exploring uncertainty, while preserving quantitative agreement with the combinatorial approach.

### 3.2 Incidence of arrhythmic events and threshold exposures

The concentration dependence of arrhythmic outcomes is summarized in Figure 4. In both saline buffer and human plasma conditions, the probability of arrhythmia (*P* (Arrhythmia)) increases sharply once total concentrations exceeded distinct, sex-dependent thresholds. Under saline buffer conditions, arrhythmia onset occurs at 943 ng/mL for females and 1569 ng/mL for males in the combinatorial predictions, and at 964 ng/mL (female) and 1693 ng/mL (male) in the Sobol-sampled predictions. When simulated in human plasma, thresholds are markedly lower—434 ng/mL (female) and 1138 ng/mL (male) for combinatorial sampling, and 427 ng/mL (female) and 849 ng/mL (male) for Sobol sampling—reflecting higher free drug exposure under plasma binding conditions.

**Figure 4:**
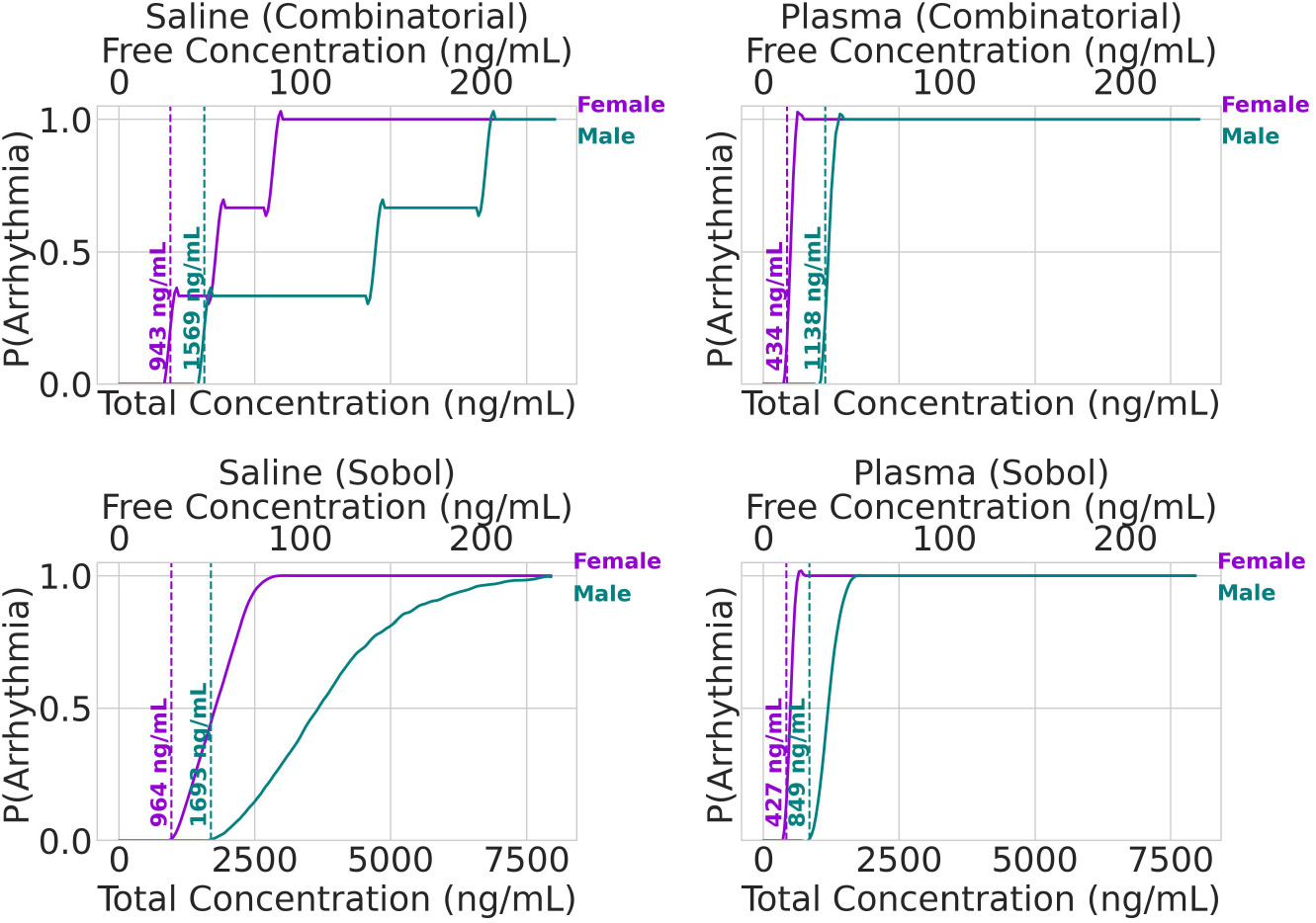
Concentration-dependent probability of arrhythmia. Probability of arrhythmia (*P* (Arrhythmia)) as a function of loperamide total plasma concentration (bound + unbound) and free plasma concentration (unbound, pharmacologically active fraction) in virtual male (teal) and female (magenta). Results are shown for saline buffer and human plasma conditions using both combinatorial (top) and Sobol (bottom) sampling. Dashed vertical lines indicate the onset thresholds at which arrhythmic events first appear.

Across all simulations, arrhythmic events are absent at therapeutic exposures and appear only at supratherapeutic concentrations. The steep rise in *P* (Arrhythmia) near the onset thresholds indicates a narrow transition between non-arrhythmic and fully arrhythmic regimes. Female models consistently exhibit lower arrhythmogenic thresholds than male models, underscoring increased susceptibility to loperamide-induced arrhythmias. Once arrhythmia probability reaches unity, the corresponding plateau regions occur at concentrations far exceeding therapeutic levels, highlighting that clinically relevant arrhythmogenic risk arises only under conditions of severe overexposure.

Importantly, near the arrhythmia onset thresholds, ΔQT variability is maximal (see Figures 2 and 3), with some predictions exhibiting modest prolongation while others develop overt arrhythmias at the same exposure level. These findings underscore that risk cannot be defined solely by mean behavior but emerges from the interaction of concentration, biological variability, and sex-specific factors.

Although these concentrations span a wide range that ensures robust testing of the emulators’ performance, they do not directly align with the clinical literature. Case reports have documented arrhythmic events at substantially lower plasma concentrations (Marraffa et al., 2014; Modi et al., 2021), typically between 22 and 130 ng/mL. Nevertheless, several studies emphasize that these reported values may not accurately reflect true systemic exposure and are therefore an unreliable predictor of adverse events (Mukarram et al., 2016).

### 3.3 Targeted emulators’ validation under extreme conditions

To evaluate whether the emulators could replicate the predictions of the simulator, a head-to-head comparison at selected exposure levels was performed (Tables 2) using saline buffer data. Three representative concentrations were chosen, spanning non-arrhythmic (82 ng/mL), borderline arrhythmic (1999 ng/mL), and overtly arrhythmic (4033 ng/mL) conditions. Across these scenarios, emulators’ predictions of ΔQT closely match those of the simulator, with both models identifying the same arrhythmic outcomes. Small discrepancies in ΔQT are observed, with differences increasing as concentration rises and emulators’ reliability decreases, and only a single mismatch is noted at the arrhythmic onset concentration.

**Table 2:** Emulators’ vs simulator’s predictions with saline buffer data. ΔQT is not computable when arrhythmia is predicted.

| Conc. (ng/mL) | Female |  | Male |  |
| --- | --- | --- | --- | --- |
| | Emulator<br>(ΔQT/Arrhythmia) | Simulator<br>(ΔQT/Arrhythmia) | Emulator<br>(ΔQT/Arrhythmia) | Simulator<br>(ΔQT/Arrhythmia) |
| 82 | 2.34 ms / No | 0 ms / No | 2.21 ms / No | 4 ms / No |
| 82 | 11.29 ms / No | 8 ms / No | 10.30 ms / No | 12 ms / No |
| 1999 | - / Yes | - / Yes | 157.98 ms / No | 148 ms / No |
| 1999 | - / Yes | - / Yes | - / Yes | 280 ms / No |
| 4033 | - / Yes | - / Yes | 180.41 ms / No | 172 ms / No |
| 4033 | - / Yes | - / Yes | - / Yes | - / Yes |

## 4 Discussion

This study demonstrates that computational cardiac emulators can extend proarrhythmic risk assessment into domains that are typically inaccessible to experimental testing and even to traditional mechanistic modeling: the explicit propagation of input variability and the systematic exploration of overdose conditions. Using loperamide as a test case, we showed that variability in ion-channel measurements substantially broadens predicted outcomes, while extreme exposure simulations reveal sex-specific thresholds for arrhythmic onset. Furthermore, the adoption of emulators offers a substantial computational advantage compared with traditional 3D electrophysiology simulations. In practical terms, reproducing a single case with the simulator takes on average 3.43 hours, whereas the emulators can predict ΔQT in mere centiseconds, achieving a speed-up of roughly five orders of magnitude. Together, these findings highlight both the power and the necessity of variability-aware, sex-specific in silico frameworks in modern safety pharmacology.

A key observation is that experimental uncertainty, particularly in IC_50_ and Hill slope parameters, strongly influenced the spread of predicted ΔQT and arrhythmia incidence. This variability was most pronounced near arrhythmic threshold concentrations, where small changes in input data led to divergent outcomes ranging from modest ΔQT values to overt arrhythmias. Such findings underscore that risk cannot be meaningfully captured by single mean values alone, supporting the idea of integrating experimental variability into safety assessment pipelines. Importantly, the correction for functional plasma protein binding narrowed the prediction envelope, suggesting that saline-based assays may exaggerate apparent uncertainty relative to more physiologically relevant conditions, such as human plasma.

The overdose simulations revealed arrhythmogenic thresholds of approximately 107 *−* 109×*C*_max_ in female models and 213 *−* 286×*C*_max_ in male models under plasma-adjusted conditions, consistent with the clinical observation that females are more susceptible to drug-induced QT prolongation. These thresholds broadly align with reported loperamide intoxication cases, where arrhythmias have been documented at supratherapeutic plasma levels (Marraffa et al., 2014; Modi et al., 2021), albeit with poor correlation between serum concentration and clinical outcome (Mukarram et al., 2016). This discrepancy likely reflects the multifactorial nature of loperamide-induced arrhythmia: affected individuals may have preexisting cardiac conditions, abuse other substances (e.g., alcohol or opioids), experience electrolyte imbalances, or encounter physiological stressors, all of which can independently lower the threshold for arrhythmic events. Collectively, these factors obscure the relationship between serum concentrations and clinical outcomes.

Another important outcome is the demonstration that machine learning emulators can replicate the predictions of full biophysical simulators with high fidelity across both safe and arrhythmogenic regimes. This scalability is critical, as exhaustive exposure analyses would otherwise be computationally prohibitive. By enabling rapid and reproducible large-scale testing, emulators open the door to systematic, variability-aware preclinical assessments that can complement traditional assays and guide regulatory decision-making.

Overall, the present study demonstrates the translational potential of computational emulators in safety pharmacology. For regulators, such frameworks could provide a rapid means of quantifying the range of possible outcomes under uncertainty, rather than relying solely on point estimates. For pharmaceutical development, they offer a tool to explore safety margins under extreme exposures or in vulnerable subpopulations before moving into costly or ethically constrained studies. Future work should aim to integrate pharmacokinetic models, expand to additional sources of biological variability, and validate emulator-based predictions against large-scale clinical and pharmacovigilance datasets.

## 5 Limitations

This work also points some limitations. First, the framework depends on the quality and completeness of the input ion-channel data; the lack of direct plasma-based measurements for non-hERG channels required extrapolation, which may introduce bias. Second, while sex-specific ion channel behavior was incorporated, other sources of variability—such as age, comorbidities, or chronic drug use—were not addressed. Third, plasma concentrations remain an imperfect surrogate for tissue exposure, particularly in cases of loperamide where gastrointestinal and distributional effects dominate toxicokinetics. Fourth, although the initial approach employed explored variability through exhaustive combinatorial pairing of available IC_50_ and Hill values, more formal uncertainty quantification approaches such as Sobol sequences can capture parameter interactions more systematically and efficiently, and may be a preferred methodology in future work. Finally, emulators’ outputs are constrained by the assumptions embedded in the underlying biophysical models, including the choice of cellular dynamics and drug–channel interaction schemes.

## 6 Conclusions

This study illustrates how sex-specific cardiac emulators can be used to efficiently propagate experimental variability and assess arrhythmic risk at exposures far beyond therapeutic ranges. In the case of loperamide, this framework captured the widening effect of assay variability on predicted outcomes and identified sex-dependent thresholds for arrhythmia onset at extreme doses. These results demonstrate that variability-aware computational emulators can complement conventional assays by providing rapid mechanistic insight into scenarios that are difficult to probe experimentally, including misuse and overdose. Such approaches have the potential to improve translational safety assessment, guide regulatory evaluation, and ultimately reduce the risk of unforeseen proarrhythmic liabilities.

## 7 Additional Information

### Author Contributions

We would like to acknowledge the significant contributions made by all authors to this work. P.D.-G. contributed to Conceptualization, Data Curation, Formal Analysis, Investigation, Methodology, Project Administration, Software, Validation, Visualization, Writing – Original Draft, and Writing – Review & Editing. A.Z. and C.B. contributed to Software, Supervision, Validation, Writing – Original Draft, and Writing – Review & Editing. M.L., N.J.-P. and G.R. contributed to Conceptualization, Data Curation, Methodology, Resources, Writing – Original Draft, and Writing – Review & Editing. M.V. contributed to Funding Acquisition, Project Administration, Resources, Software, Supervision, and Writing – Review & Editing. J.A. contributed to Conceptualization, Investigation, Methodology, Project Administration, Software, Supervision, Validation, Writing – Original Draft, and Writing – Review & Editing.

All authors approved the final version of the manuscript and agree to be accountable for all aspects of the work, ensuring that any questions related to the accuracy or integrity of the work are appropriately investigated and resolved. All authors meet the criteria for authorship and are listed accordingly.

### Data Availability

ELEM Biotech owns the commercial rights to Alya, the computational finite element solver employed to train the emulators. The methodology can be replicated using any finite element solver given all the models’ information provided in this paper and previously published literature.

The data generated and analyzed during this study are proprietary to ELEM Biotech and cannot be publicly shared due to commercial confidentiality. However, access to the data may be considered on a case-by-case basis. Requests for access can be directed to ELEM Biotech compliance department, and will be subject to a data use agreement ensuring compliance with relevant regulations and restrictions.

